# WRN Helicase is a Synthetic Lethal Target in Microsatellite Unstable Cancers

**DOI:** 10.1101/502070

**Authors:** Edmond M. Chan, Tsukasa Shibue, James McFarland, Benjamin Gaeta, Justine S. McPartlan, Mahmoud Ghandi, Jie Bin Liu, Jean-Bernard Lazaro, Nancy Dumont, Alfredo Gonzalez, Annie Apffel, Syed O. Ali, Lisa Leung, Emma A. Roberts, Elizaveta Reznichenko, Mirazul Islam, Maria Alimova, Monica Schenone, Yosef Maruvka, Yang Liu, Alan D’Andrea, David E. Root, Jesse S. Boehm, Gad Getz, Todd R. Golub, Aviad Tsherniak, Francisca Vazquez, Adam J. Bass

## Abstract

Synthetic lethality, an interaction whereby the co-occurrence of two or more genetic events lead to cell death but one event alone does not, can be exploited to develop novel cancer therapeutics^1^. DNA repair processes represent attractive synthetic lethal targets since many cancers exhibit an impaired DNA repair pathway, which can lead these cells to become dependent on specific repair proteins^2^. The success of poly (ADP-ribose) polymerase 1 (PARP-1) inhibitors in homologous recombination-deficient cancers highlights the potential of this approach in clinical oncology^3,4^. Hypothesizing that other DNA repair defects would give rise to alternative synthetic lethal relationships, we asked if there are specific dependencies in cancers with microsatellite instability (MSI), which results from impaired DNA mismatch repair (MMR). Here we analyzed data from large-scale CRISPR/Cas9 knockout and RNA interference (RNAi) silencing screens and found that the RecQ DNA helicase *WRN* was selectively essential in MSI cell lines, yet dispensable in microsatellite stable (MSS) cell lines. WRN depletion induced double-strand DNA breaks and promoted apoptosis and cell cycle arrest selectively in MSI models. MSI cancer models specifically required the helicase activity, but not the exonuclease activity of WRN. These findings expose *WRN* as a synthetic lethal vulnerability and promising drug target in MSI cancers.

## Main Text

Defects of DNA mismatch repair (MMR) promote a hypermutable state characterized by frequent insertion/deletion mutations, which occur preferentially at nucleotide repeat regions known as microsatellites, and single-nucleotide variant (SNV) mutations^5^. This class of hypermutation, termed microsatellite instability (MSI), contributes to the development of several cancers, including 15% of colon cancers^7^, 22% of gastric cancers^8^, 20–30% of endometrial cancers^9,10^, and 12% of ovarian cancers^11^. MSI cancers can arise in the setting of germline mutations in the MMR genes *MSH2, MSH6, PMS2*, or *MLH1*, a condition known as Lynch Syndrome^5^. More commonly, MSI cancers develop following somatic inactivation of an MMR gene, typically *MLH1* loss via promoter hypermethylation^5^. Recently, MSI cancers have been associated with significant responses to immune checkpoint inhibition. However, 45–60% of patients with MSI cancers do not respond to immune checkpoint blockade and the use of these agents can also be limited by their toxicity^12,13^. Hence, there remains a significant need to develop further therapies targeting MSI tumors.

We hypothesized that MMR deficiency may create unique vulnerabilities in MSI cancers. To evaluate this hypothesis and identify candidate therapeutic targets for MSI cancers, we analyzed two independent large-scale cancer dependency datasets, Project Achilles and Project DRIVE. The Project Achilles dataset consisted of 391 cancer cell lines screened with a near genome-wide CRISPR/Cas9 library and Project DRIVE interrogated 398 cancer cell lines with an RNAi library to define genes essential for proliferation and survival of individual cancer cell lines^14,15^. By comparing essential genes in cell lines with or without specific features such as MSI, these functional genomic datasets provide an opportunity to identify synthetic lethal interactions (Fig. 1a). Moreover, the large number of cell lines included in these screens helps to more robustly identify dependencies associated with distinct features of cancer and not merely idiosyncratic dependencies isolated to individual cell lines.

**Fig. 1.**
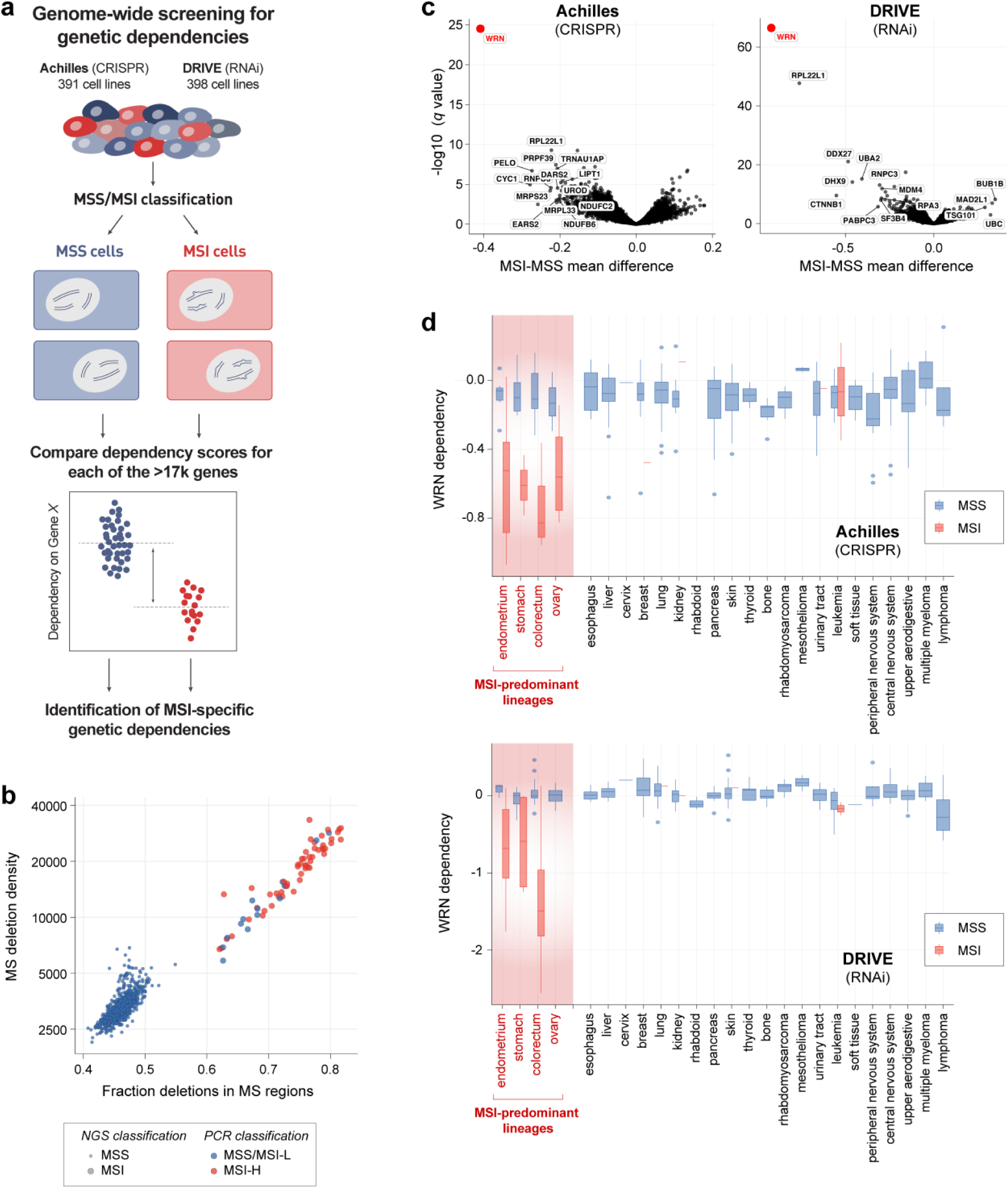
Genome-wide loss-of-function screening identifies genes synthetic lethal with MSI. **a**, Schematic of dependency dataset analysis. Cell lines were grouped by feature and dependency scores for each gene were analyzed to identify feature-specific genetic dependencies. **b**, Screened cell lines were plotted by the deletion density and fraction of deletions occurring in microsatellite (MS) regions. MSI classification by next generation sequencing (NGS) and multiplex polymerase chain reaction (PCR) are indicated. **c**, False discovery rate (FDR) q-values were plotted against the mean difference of dependency Z scores between MSI and MSS cell lines for both Project Achilles CRISPR/Cas9 and Project DRIVE RNAi datasets. **d**, *WRN* dependency scores were plotted by cancer lineage and further subclassified by MSI and MSS status. Gene dependency scores are normalized to the control sgRNAs for each cell line. A value 0 represents the median dependency score of negative control sgRNAs and −1 represents the median dependency score of sgRNAs targeting pan-essential control genes. The lower and upper hinges correspond to first and third quartiles (25^th^ and 75^th^ percentiles). The upper and lower whiskers extend to the largest value within 1.5*IQR (inter-quartile range) from the hinge.

Before we could test this approach, we needed to differentiate cell lines with MSI from microsatellite stable (MSS) models. The clinical MSI assay is performed by a single multiplex polymerase chain reaction (PCR) and analyzed by capillary electrophoresis^16^. However, these data were not available for all screened cell lines. Instead, we utilized MSI classifications from Phase II of the Cancer Cell Line Encyclopedia (CCLE) project^17^, which analyzed next-generation sequencing (NGS) data to determine the total number of genetic deletions and fraction of deletions located within microsatellite regions. These analyses, which were reproduced to facilitate comparison with dependency data, identified two distinct groups that were characterized as MSI and MSS (Fig. 1b). Moreover, these MSI classifications were highly concordant with available PCR-based MSI phenotyping^18,19^ and with the predicted presence of MMR loss as defined by loss of mRNA expression, deletion, and/or damaging mutations to *MLH1, MSH2, MSH6*, and/or *PMS2*. In total, 45 unique MSI and 480 unique MSS cell lines were represented by one or both datasets.

We then compared genetic dependencies between MSI and MSS cell lines in each screening dataset. The Project Achilles and Project DRIVE datasets both independently identified *WRN*, which encodes a RecQ DNA helicase, to be the top preferential dependency in MSI cell lines (q-value 5.3×10^−21^ and 2.6×10^−63^, respectively, Fig. 1c). These findings remained true when we evaluated the data using the PCR-based MSI classifications. In contrast, none of the four other RecQ DNA helicases were identified as a preferential dependency in MSI cell lines.

MSI is most commonly observed in colorectal, endometrial, gastric, and ovarian cancers, which were well-represented by cell lines in the screening datasets. Indeed, when these data were stratified by lineage and MSI status, MSI cell lines from these four lineages (n = 34) showed greater dependence upon *WRN* than their MSS counterparts (n = 130). We also identified 11 MSI cell lines from lineages where MSI is less common (4 leukemia, 2 prostate, and single models of other lineages). However, these MSI cell lines were phenotypically distinct from the MSI cell lines of the four MSI-prone lineages. These ‘atypical’ MSI models were both less dependent on *WRN* (Fig. 1d) and possessed a substantially lower density of deletion mutations in microsatellite regions, despite also possessing events predictive of MMR inactivation. These observations raised the possibility that *WRN* dependency is not simply a result of MMR deficiency, but it may require specific cancer lineages and/or a more profound mutator phenotype. Indeed, even within the MSI cell lines from the four MSI-predominant lineages, *WRN* dependency correlated with the density of microsatellite deletions (Spearman’s rho = −0.54, *p* = 0.0012).

These data suggested that *WRN* is a selective dependency in MSI cancers, raising the hypothesis that WRN could be a novel drug target for this class of cancers. To assess *WRN* as a genetic dependency, we validated three sgRNAs targeting *WRN* by demonstrating depletion of the WRN protein by immunoblot (IB) (Fig. 2a). We then evaluated the effects of CRISPR/Cas9-mediated *WRN* knockout in 5 MSS and 5 MSI cell lines, all from the four MSI-predominant lineages, with an 8-day viability assay. The effects of *WRN* knockout were comparable to the positive ‘pan-essential’ controls in the MSI lines. By contrast, *WRN* silencing in MSS models approximated the effects of negative controls, targeting intergenic regions (Fig. 2b). Similarly, in a 10-day competitive growth assay, CRISPR/Cas9-mediated WRN depletion substantially impaired the viability of MSI cell lines despite negligible effects in MSS cell lines (Fig. 2c). Complementing the CRISPR/Cas9 data, RNAi-mediated *WRN* silencing with short hairpin RNA (shRNA) impaired MSI, but not MSS, cell viability (Figs. 2d, 2e), consistent with our hypothesis that WRN loss is synthetic lethal with MSI.

**Fig. 2.**
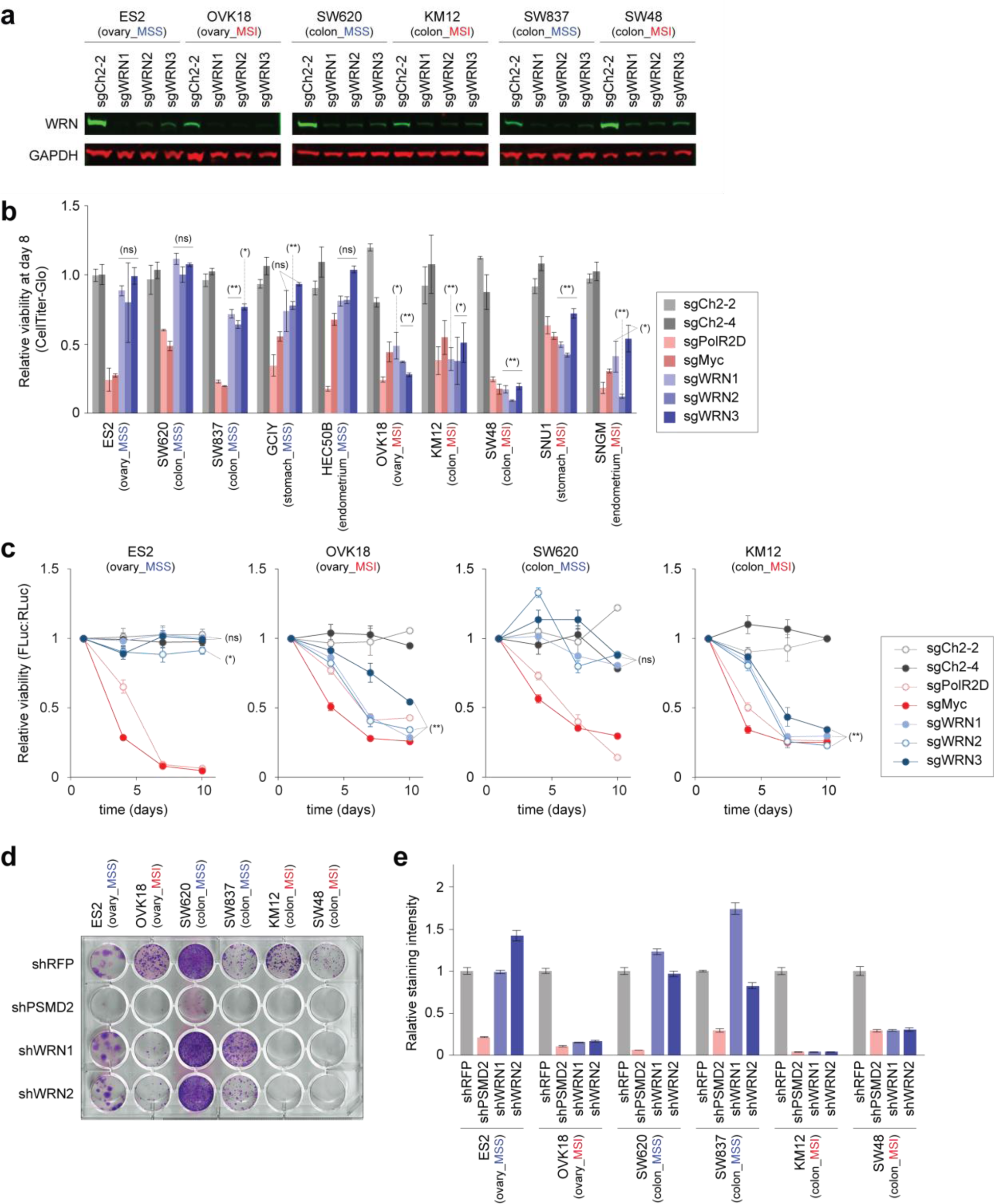
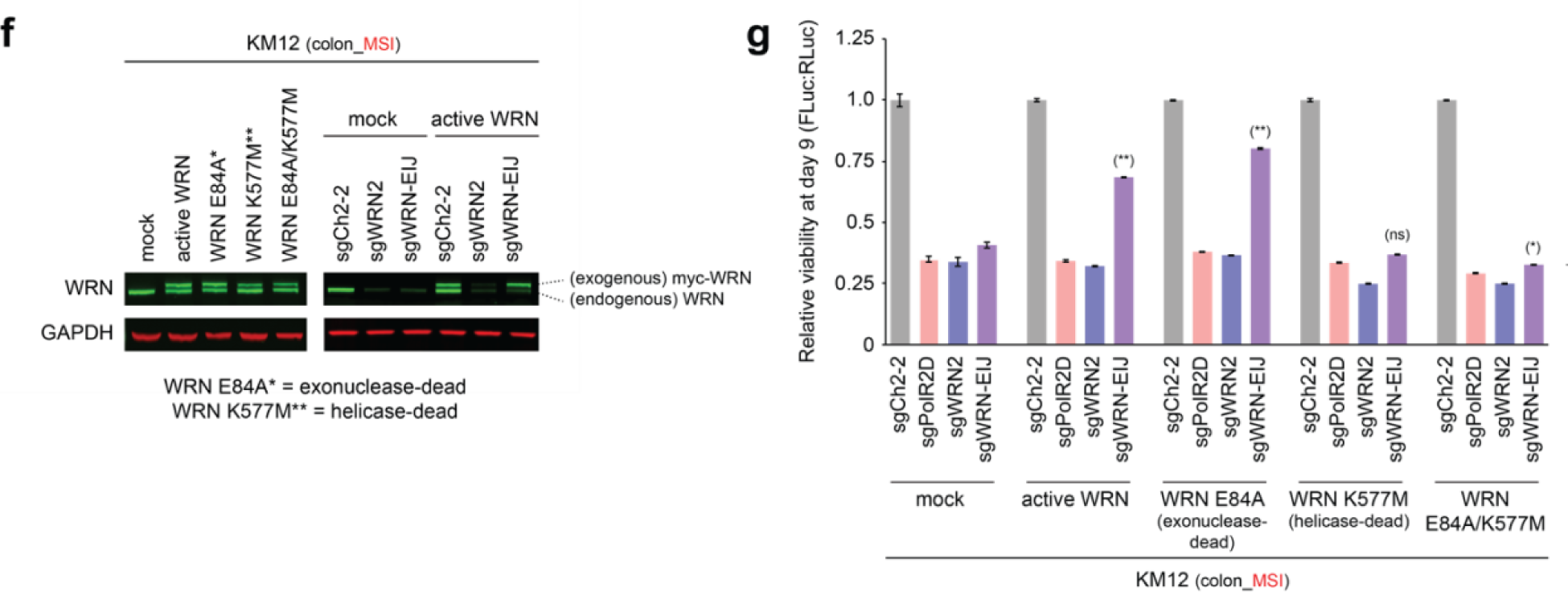
WRN is a synthetic lethal partner with MSI. **a**, Immunoblot (IB) of WRN and GAPDH (loading control) in representative Cas9-expressing MSS and MSI cell lines 4 days after lentiviral transduction with indicated sgRNAs. **b**, Relative viability of Cas9-expressing cell lines 8 days after lentiviral delivery of the indicated sgRNAs: negative controls targeting intergenic sites of chromosome 2 (sgCh2–2, sgCh2–4), pan-essential controls (sgPolR2D, sgMyc) and *WRN* (sgWRN1, sgWRN2, sgWRN3). Values = mean ± SE (n = 3 [b], n = 6 [c]). **c**, Differential cell viability of representative MSS and MSI cell lines treated with indicated sgRNAs in a luciferase-based competitive growth assay. **d**, Clonogenic assay of indicated cell lines after lentiviral transduction with the indicated shRNAs: non-targeting negative control (shRFP), pan-essential control (shPSMD2), and 2 shRNAs targeting *WRN* (shWRN1, shWRN2). **e**, Relative staining intensity of the clonogenic assay. **f**, (left) IB of KM12 cells expressing myc-tagged active, exonuclease-dead (E84A), helicase-dead (K577M), or dually exonuclease/helicase-dead (E84A/M577M) versions of *WRN* cDNA. (right) IB of cells transduced with sgRNAs targeting an *WRN* exon (sgWRN2) or exon-intron junction (sgWRN-EIJ). **g**, Luciferase-based competitive growth assay 9 days post-transduction of indicated sgRNAs in KM12 cells expressing GFP or indicated version of WRN. Values are presented as means ± SE (n = 6). (*) p < 0.05; (**) p < 0.001; (ns) p ≥ 0.05 by two-tailed Student’s *t*-test.

To test whether the phenotype elicited by the sgRNAs was specifically due to inactivation of WRN, we developed an sgRNA that binds to an exon-intron junction of *WRN* (sgWRN-EIJ) and hence would target endogenous *WRN* but not exogenous *WRN* cDNA. Cas9-expressing KM12 cells, a MSI colorectal cancer model, were then transduced with *GFP* or myc-tagged *WRN* cDNA followed by distinct sgRNAs. Indeed, sgWRN-EIJ silenced endogenous *WRN* but not exogenous *WRN* cDNA (Fig. 2f). Correspondingly, *WRN* cDNA but not GFP expression rescued KM12 cell viability from sgWRN-EIJ (Fig. 2g). In contrast, sgWRN2, targeting both endogenous and exogenous *WRN*, equally impaired both GFP-expressing and *WRN* cDNA-expressing cell viability. These observations argue that the selective lethal phenotype induced by sgRNAs targeting *WRN* is attributable to its effects upon *WRN* and not an ‘off-target’ effect.

Together, these findings suggest that the WRN-dependent state in MSI/MMR-deficient cancers could be therapeutically exploited by targeted inhibition of WRN. The WRN protein functions as both a 3’ to 5’ exonuclease and 3’ to 5’ helicase in multiple processes including DNA repair, DNA replication, and telomere maintenance^20^. Thus, we sought to determine which enzymatic function of WRN is essential for MSI cancer cell lines. Using our ability to reintroduce *WRN* cDNA to rescue the effects of targeting endogenous *WRN*, we expressed exonuclease-dead (E84A), helicase-dead (K577M), or dually exonuclease/helicase-dead (E84A/K577M) versions of *WRN* cDNA^20^ in KM12. We found that inactivation of the exonuclease domain did not attenuate rescue, suggesting this enzymatic function of WRN is dispensable in MSI cancers. By contrast, inactivation of the WRN helicase prevented rescue (Fig. 2g). By replicating the dependency of MSI cancers on the WRN protein with genetic inhibition of the helicase function of WRN, these results nominate the helicase domain as a candidate drug target.

We next sought to understand the cellular consequences of WRN loss, using both CRISPR/Cas9 and shRNA-mediated silencing of *WRN* in MSI and lineage-matched MSS models. Cell cycle analysis with EdU/DAPI staining revealed that *WRN* silencing reduced the proportion of MSI cells in S phase. Concomitantly, we observed an increase in cells in G1 or G2/M phases, suggesting cell cycle arrest at either the G1 or G2/M phases (Fig. 3a). Furthermore, annexin V/propidium iodide (PI) staining demonstrated induction of apoptosis and evidence of cell death in MSI cells following *WRN* silencing (Fig. 3b). In contrast, MSS cell lines demonstrated no discernable evidence of cell cycle arrest nor increased cell death after *WRN* silencing.

**Fig. 3.**
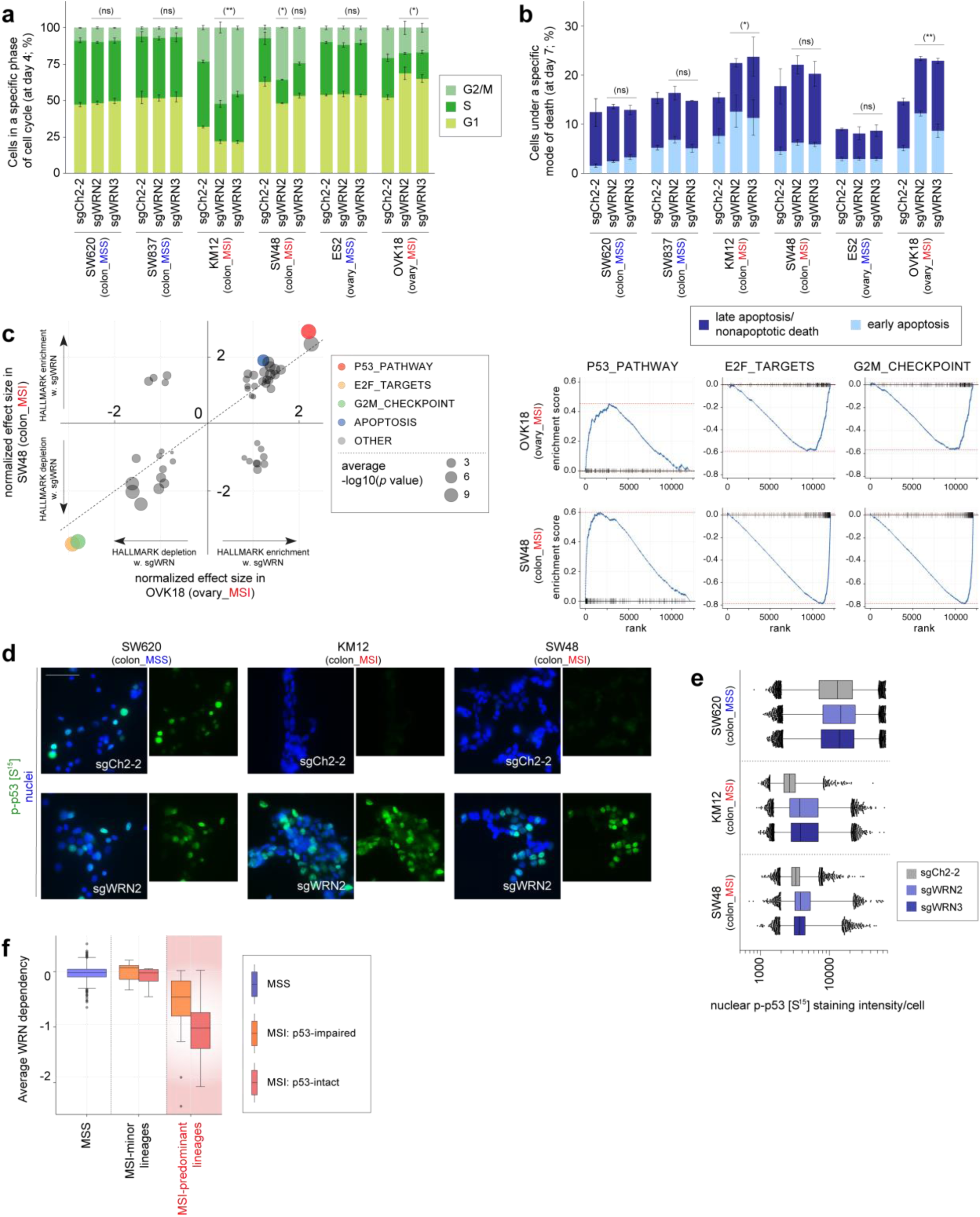
WRN depletion in MSI cells induces cell cycle arrest, apoptosis, and a p53 response. **a**, Cell cycle evaluation of representative Cas9-expressing MSI and MSS cell lines after lentiviral transduction with the indicated sgRNAs. **b**, Annexin V/propidium iodide (PI) staining evaluating early apoptosis and late apoptosis/non-apoptotic cell death in representative Cas9-expressing MSI and MSS cell lines 7 days after delivery of indicated sgRNAs. **c**, GSEA enrichment and depletion scores in WRN-depleted OVK18 cells plotted against WRN-depleted SW48 cells. Signature enrichment plots for the indicated Hallmark gene sets are shown for OVK18 and SW48 cells after WRN depletion. **d**, phospho-p53 (S15) immunofluorescence (IF) of indicated Cas9-expressing cell lines treated with indicated sgRNAs. Scale bar = 50 μm. **e**, Quantification of nuclear phospho-p53 (S15) staining intensity per cell. **f**, *WRN* dependency was evaluated for cell lines classified as MSS, MSI from an infrequent MSI lineage, or MSI from an MSI-predominant lineage and further subclassified by p53 status. (*) p < 0.05; (**) p < 0.005; (ns) p ≥ 0.05 by two-tailed Student’s *t*-test. Hinges and whiskers as per Fig. 1d.

We next pursued the molecular basis for the dependence upon *WRN* in MSI cell lines. Consistent with the aforementioned cell cycle and apoptosis assays, evaluation of the transcriptional effects of *WRN* silencing in MSI cells with mRNA sequencing followed by gene set enrichment analysis (GSEA) demonstrated upregulation of the apoptotic signature and downregulation of G2/M checkpoint progression signature. This analysis also highlighted the activation of p53 as the most upregulated transcriptional signature following WRN loss in MSI cells (Fig. 3c). Further investigation with immunofluorescence (IF) in the MSI models after *WRN* silencing demonstrated increased p53 phosphorylation at S15, a key signal for p53 activation^21^. In contrast, we observed no substantial change in the phospho-p53 signal intensity in the MSS models after WRN depletion (Figs. 3d, 3e). After *WRN* depletion in the *TP53* wild-type MSI cells, we also observed increased staining of cyclin-dependent kinase inhibitor p21, providing another indication of p53 activation. Conversely, we observed no substantial change in p21 signal intensity after *WRN* silencing in our MSS and TP53-mutant MSI cells. We then re-evaluated our dependency data, stratifying by MSI and p53 status. While p53-intact MSI cell lines were more sensitive to WRN loss than their p53-impaired counterparts, this analysis demonstrated that both wild-type and mutant *TP53* MSI cell lines were dependent on *WRN* (Fig. 3f). These data suggest that while WRN loss leads to p53 activation, p53 activity is not solely responsible for WRN dependence.

The finding of increased p53 phosphorylation at S15, a phosphorylation target of the DNA-damage response kinases ATR and ATM^21^, and subsequent p53 activation led us to hypothesize that WRN loss in MSI cancer cells may lead to DNA damage. This hypothesis is consistent with the well-reported roles of p53 and WRN in responding to DNA damage and preserving DNA integrity^22–24^. Indeed, biallelic germline inactivation of *WRN* leads to Werner Syndrome, a disease characterized by premature aging and increased cancer incidence due to impaired DNA damage repair and telomeric shortening leading to chromosomal aberrations^25,26^. We therefore asked if DNA damage may be responsible for the impaired viability observed in WRN-depleted MSI cells. Consistent with this hypothesis, *WRN* silencing in MSI cells, but not MSS cells, substantially increased γH2AX and 53BP1 foci, markers of double-strand DNA breaks (DSB) (Figs. 4a-c). These findings were further corroborated by increased phospho-ATM (S1981) foci formation and Chk2 phosphorylation at T68, indicating a cellular DSB response known to activate p53 and anti-proliferative signaling^27^ (Figs. 4a, 4d). We also observed increased γH2AX in MSI cells transduced with shRNA against *WRN*, arguing that the DSBs are not just a consequence of CRISPR/Cas9 endonuclease activity (Fig. 4e). These observations explain why p53-impaired MSI cells are still sensitive to WRN depletion since DSBs are toxic to cells, independent of p53 status^28^.

**Fig. 4.**
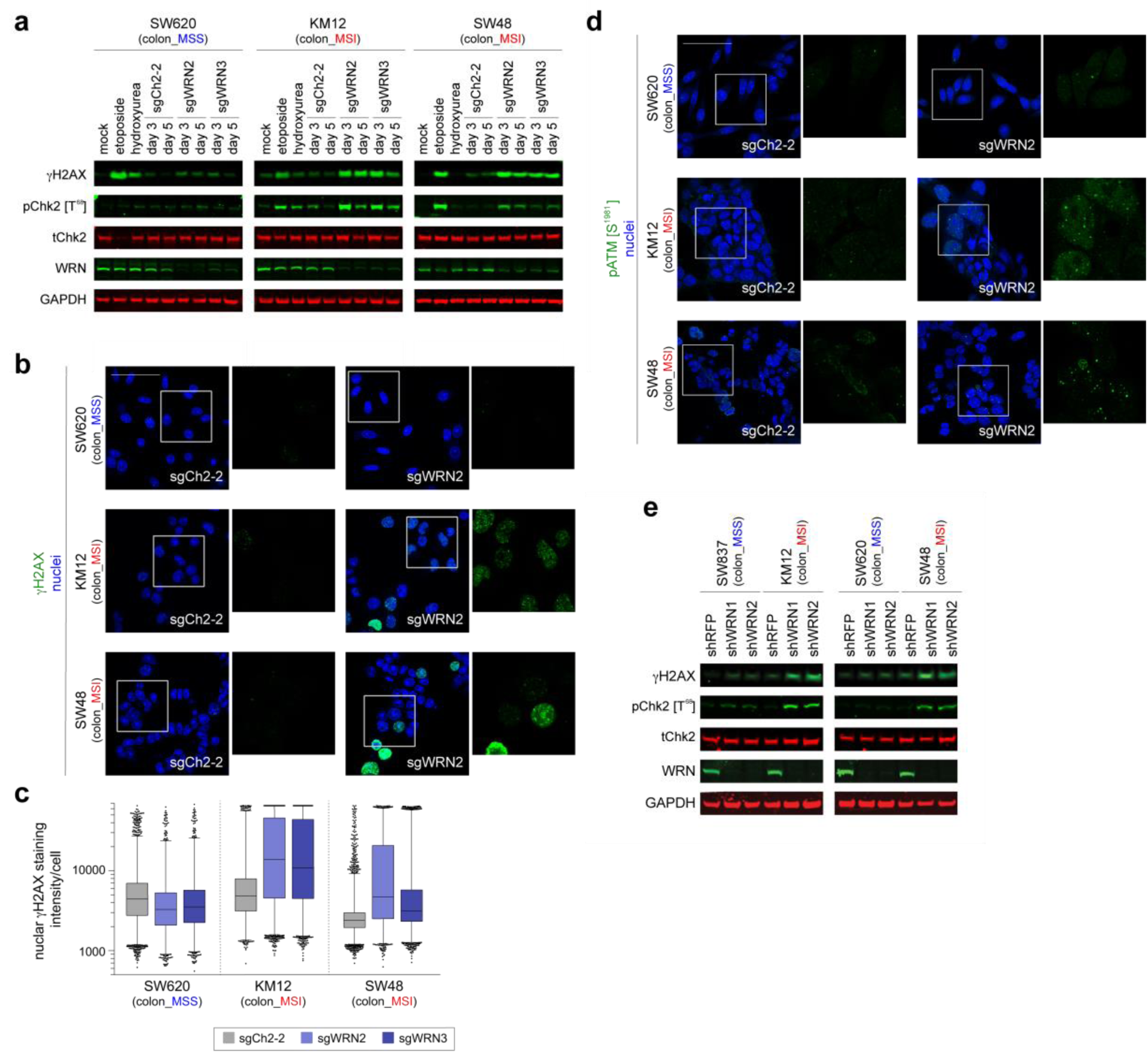
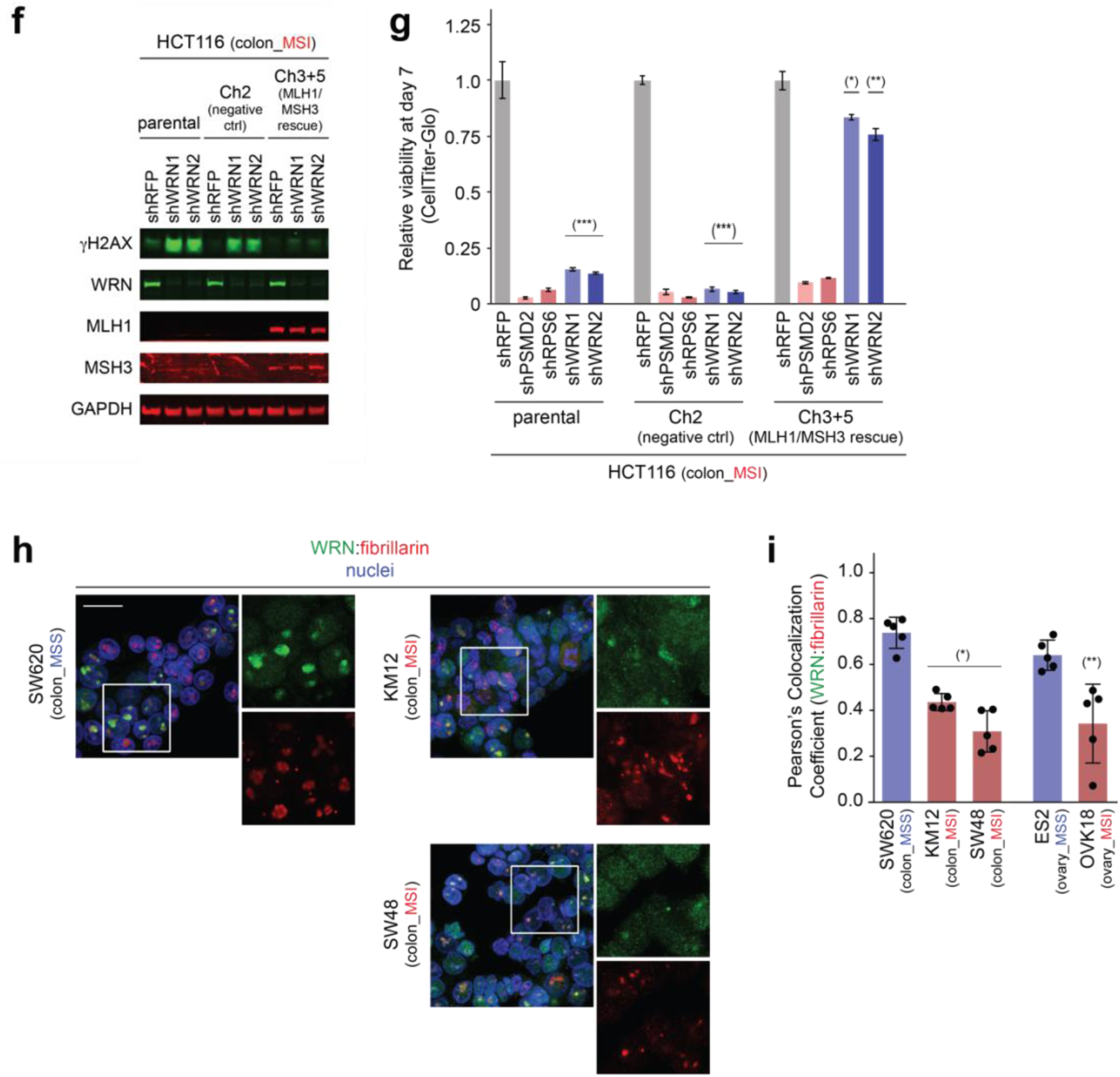
WRN depletion in MSI cells leads to accumulation of double strand DNA breaks. **a**, IB of γH2AX, phospho(T86)- and total-Chk2, WRN, and GAPDH in representative cell lines after CRISPR/Cas9-mediated WRN depletion. **b**, γH2AX IF of representative cell lines after delivery of indicated sgRNAs. Scale bar = 50 μm. **c**, Quantification of nuclear γH2AX staining intensity per cell. **d**, phospho-ATM (S1981) IF of representative cell lines after delivery of indicated sgRNAs. Scale bar = 50 μm. **e**, IB of γH2AX, phospho(T86)- and total-Chk2, and GAPDH after RNAi-mediated *WRN* knockdown. **f**, IB of γH2AX, WRN, MLH1, MSH3, GAPDH in HCT116 with or without MMR restoration after lentiviral transduction of indicated shRNAs. **g**, Relative viability of HCT116 cells with and without MMR restoration 7 days after lentiviral delivery of the indicated shRNAs. Values are presented as means ± SE (n = 6). (*) p < 0.01; (**) p < 0.001; (***) p < 0.0002 by two-tailed Student’s *t*-test. **h**, WRN IF of representative cell lines. Scale bar = 20 μm **i**, Quantitative analysis of WRN colocalization to the nucleolar marker, fibrillarin, by Pearson’s colocalization coefficients. (*) p < 0.001; (**) p < 0.02 by two-tailed Student’s *t*-test.

We next sought to determine which feature of MSI cells, the hypermutable state or MMR deficiency, leads to DNA damage and viability impairment upon WRN loss. We first considered the possibility that the hypermutation in MSI cells leads to recurrent mutation and inactivation of another helicase, creating a dependency upon the WRN helicase. Analogously, the second most significant dependency in MSI models from both datasets was *RPL22L1*, a previously defined dependency often found in MSI cancers due to frequent mutation of its paralog, RPL22^15,29^ (Fig. 1c). However, we found no other helicase whose loss could account for the preferential dependency upon *WRN* in MSI cells. We also asked if cell lines with other hypermutable states similarly require WRN. Specifically, we evaluated cell lines with polymerase epsilon *(POLE)* exonuclease domain mutations that have been described to impair the *POLE* proofreading function. These mutations induce a markedly increased frequency of SNV mutations rather than the mixed pattern of insertion/deletion and SNV mutations associated with MMR deficiency^30,31^. Using published reports of *POLE* mutations implicated to cause a hypermutator phenotype, we analyzed the functional genomic datasets and identified 5 cell lines with reported proofread-impairing *POLE* mutations. All 5 cell lines were MSS from endometrial or colorectal lineages and insensitive to WRN loss, suggesting that hypermutability alone cannot account for *WRN* dependency.

We then explored whether MMR deficiency contributes the MSI cell’s dependence upon *WRN*. To assess this hypothesis, we compared the effects of *WRN* silencing upon a previously characterized model of MMR restoration. In this model, the MMR activity of the *MLH1-* and *MSH3-mutated* MSI cell line, HCT116, was functionally restored by introducing chromosomes 3 and 5 carrying a wild-type copy of *MLH1* and *MSH3*, respectively^32,33^. *WRN* knockdown led to γH2AX accumulation and impaired the viability of both parental HCT116 and a control HCT116 cell line with an additional chromosome 2. By contrast, transfer of chromosomes 3 and 5 from normal fibroblasts suppressed γH2AX accumulation and partially rescued HCT116 viability from WRN depletion (Figs. 4f, 4g). These data suggest that ongoing MMR impairment contributes to the increased DSB and impaired viability observed in response to *WRN* silencing. One potential reason for this contextual requirement is that MMR deficiency contributes to the formation or accumulation of genomic structures that require the WRN helicase to unwind and thus prevent DNA damage. Such structures could include insertion-deletion loops^24^ and/or displacement loops (D-loops) between homeologous DNA sequences^34,35^. Indeed, MMR impairment in yeast creates a dependency on *Sgs1*, the yeast homolog of *WRN* and *BLM*, to resolve homeologous D-loops normally rejected by an intact MMR system^36^. Beyond the potential role of WRN in preventing DNA damage, loss of WRN’s capacity to participate in non-homologous end joining and/or homologous recombination (HR) could further contribute to DSB accumulation^37^.

Given the differential requirement for WRN between MSI and MSS cells, we hypothesized that WRN may have a greater role in maintaining genomic integrity following development of MSI. WRN has been reported to respond to stress upon DNA damage by disseminating from the nucleolus towards the DNA in the nucleoplasm^38,39^. We therefore compared the intracellular localization of WRN in MSI and MMR cells. IF for WRN demonstrated a predominantly dispersed pattern across the nucleoplasm in MSI cells, supporting the hypothesis that WRN is recruited to the nucleoplasm in response to stress upon the DNA. MSS cells, in contrast, demonstrated a greater degree of WRN co-localization with the nucleolar marker, fibrillarin (Figs. 4h, 4i), and less overall nucleoplasmic staining. Together with our prior observations, these data argue that WRN plays a greater and more essential role maintaining DNA integrity in MSI cells as compared to MSS cells.

Our observations have revealed that *WRN* inactivation induces DSBs and activates a DSB cellular response to promote cell death and cell cycle arrest preferentially in MSI cells. Although WRN and other DNA helicases have been nominated as possible therapeutic targets in cancer^40,41^, this work highlights WRN helicase as a synthetic lethal target in the context of MSI/MMR deficiency. As with the synthetic lethal relationship between HR deficiency and PARP inhibitor sensitivity, where 30–69% of patients with *BRCA1/2-mutant* cancers respond to PARP inhibitors^4^, *WRN* dependency in MSI cancers is not absolute. This suggests that other cellular factors mitigate this dependency. Indeed, *WRN* dependence correlated with the number of deletion mutations, raising the hypotheses that the density of microsatellite deletions reflects the degree of MMR impairment and/or mutational accumulation synergizes with MMR impairment to form DNA structures that require WRN to unwind. Nonetheless, these results support the design of WRN helicase inhibitors to pharmacologically exploit the WRN-dependent state found in MSI cancers. While systemic WRN inhibition could induce complications akin to Werner Syndrome^42^, the clinical manifestations of this syndrome require decades to emerge, suggesting that the therapeutic benefits would likely greatly outweigh these risks. Therefore, inhibitors of WRN could emerge as potent new therapy for this large and deadly class of human cancer.

## Materials and Methods

### Cancer Dependency Data

CRISPR dependency data from Project Achilles can be downloaded from the Figshare repository (https://doi.org/10.6084/m9.figshare.5863776.v1). These data contain gene dependencies estimated for each gene and cell line using the CERES algorithm^14^, which corrects for gene-independent DNA cleavage toxicity effects. RNAi dependency data were derived from Novartis’ Project DRIVE^15^, and were reprocessed using the DEMETER2 algorithm to estimate gene dependencies^43^, which can be downloaded from (https://doi.org/10.6084/m9.figshare.6025238.v1). For some analyses (Fig. 3f, Fig. S1d,e) we computed aggregate *WRN* dependency scores for each cell line by averaging together RNAi and CRISPR dependency scores (which are both normalized so that the median score of pan-essential genes is set at −1)^14,43^.

### Genomics Data

All genomic data used in the analysis has been published and can be found in the Cancer Cell Line Encyclopedia (CCLE) portal (https://portals.broadinstitute.org/ccle/data). Gene expression data (taken from the file:

CCLE_DepMap_18Q1_RNAseq_RPKM_20180214.gct) were transformed according to log(RPKM + 0.001). Gene-level relative copy number data were derived from a combination of whole-exome sequencing (WES) and SNP data to achieve maximal coverage across cell lines. When both data types were available for a given cell line, we prioritized WES over SNP data. Relative copy number data were also log-transformed for analysis. For mutation data, we utilized the merged mutation calls (file: CCLE_DepMap_18Q1_maf_20180207.txt), which combines information from multiple data sources and types.

### Cell Line Annotations

Annotations of primary disease site for each cell line can found in the DepMap portal (https://depmap.org). The functional status of TP53 in 966 cell lines was annotated based on a combination of cell lines’ nutlin-3 sensitivity (as measured in both the Genomics of Drug Sensitivity in Cancer (GDSC - http://www.cancerrxgene.org) and Cancer Target Discovery and Development (CTD2 - https://ocg.cancer.gov/programs/ctd2/data-portal) datasets), along with a p53 target gene expression signature computed from CCLE data^44^.

### Microsatellite classification

MSI classifications were obtained from Phase II of the Cancer Cell Line Encyclopedia (CCLE) project^17^. These classifications were based on the total number of deletions detected in each cell line, and the fraction of deletions in microsatellite (MS) regions, using several different data sources (CCLE WES, CCLE WGS, CCLE hybrid capture, and Sanger WES). These features were then used to classify each cell line as MSI, MSS, or indeterminate. Unless otherwise indicated, our analysis excluded cell lines classified as ‘indeterminate’. When plotting the deletion density and fraction of deletions in MS regions (Figs. 1B, S1A), we averaged these values across the data sources available for each cell line after normalizing for systematic differences between data sources. In particular, we used linear regression models to estimate and remove scale and offset differences between data sources so that the normalized number of deletions (and number of deletions in MS regions) measured in each data source was equal on average.

### MMR Status

MMR deficiency status was determined based on -omics data for the genes *MSH2, MSH6, MLH1*, and *PMS2*. For each gene we determined whether it was mutated (any detected mutation classified as either deleterious or missense), deleted (relative log2 copy number < −1), or lowly expressed (log2 RPKM mRNA expression < −1). A gene was classified as inactivated if any of the above criteria were met, and cell lines where any of these MMR genes were inactivated were classified as having ‘MMR loss’.

### Polymerase epsilon *(POLE)* exonuclease domain mutational classifications

*POLE* exonuclease domain mutational classifications were determined by compiling the previously described functional *POLE* exonuclease domain mutations from these reports^30,31,45^. Specifically, the following were considered proofread-impairing *POLE* exonuclease domain mutations: p.P286R/H/S, p.D287E, p.S297F/Y, p.V411L, p.N336S, p.P436R, p.M444K, p.A456P, pR446W, p.S459R, and p.A465V.

### Differential dependency analysis

Genes that were preferentially dependent in MSI compared to MSS cell lines were identified using linear modeling performed in parallel across genes using the R package Limma^46^. We estimated the difference in mean dependency between MSS and MSI cell lines for each gene, and associated p-values were derived from empirical-Bayes moderated t-statistics. Q-values were computed using the Benjamini-Hochberg method^47^.

### mRNA-sequencing

Cas9-expressing cells (KM12, SW48, and OVK18) were lentivirally transduced with the following sgRNAs: sgCh2-2, sgWRN2, and sgWRN3 (sequences provided below). Cells were selected with puromycin to ensure delivery of sgRNAs and RNA was purified 72 hours after transduction. This was performed in duplicate prior to cDNA library preparation and subsequent RNA-sequencing via the Illumina NextSeq 500.

We first excluded genes which had less than 1 counts per million in more than half of the samples. The weighted trimmed mean of M-values^48^ method was used to normalize the library size of each sample, using the calcNormFactors function from the R package: edgeR^49^. To estimate the log-fold change (LFC) effect of WRN knockout on each gene in each cell line we used the R package Limma^46^. Specifically, we fit a linear model for the expression of each gene, using cell line and sgRNA (WRN vs. control) as covariates. Read counts data were transformed using the Limma function ‘voom’ prior to model fitting, in order to model the mean-variance relationship of the log-counts data^50^. We then extracted LFC effect sizes and empirical-Bayes moderated t-statistics for the WRN knockout effect for each gene and cell line. Gene set enrichment analysis (GSEA)^51^ was run to test for gene sets that were up- or down-regulated in each cell line after WRN knockout. In particular, we used the R package fgsea^52^ to estimate normalized enrichment statistics, and associated p-values, for each gene set in the Hallmark Collection from the Molecular Signatures Database^53^. The GSEA algorithm was run using t-statistics as the gene-level statistics, 1 million random permutations for each cell line tested, and a “GSEA parameter” of 1.

### Cell Line and Reagents

With the exception of HCT116 and its derivatives, we obtained all the cell lines from our Cancer Cell line Encyclopedia collection^17^ HCT116 and its various derivatives were generously provided by Drs. Richard Boland, Ajay Goel and Minoru Koi^32,33^. All cell lines were grown in media supplemented with 10% fetal bovine serum (FBS), penicillin (100 μg/mL) / streptomycin (100 μg/mL) / L-glutamine (292 μg/mL; Gibco) unless otherwise stated. KM12, SW48, SW837, ES2, EFE-184 and SNU-1 were cultured in RPMI-1640 (Gibco); OVK18 were cultured in MEMα; GCIY was cultured in MEMα supplemented with 15% FBS; SW620 were cultured in Leibovitz’s L-15 (Gibco), SNGM were cultured in Ham’s F12 with 20% FBS (Gibco), HCT116 were cultured in McCoy’s 5A (Gibco). Stable Streptococcus pyogenes Cas9-expressing cell lines generated by lentiviral transduction of the pXPR_BRD111 construct were from Project Achilles^14^.

### Generation of ectopic WRN-expressing cell lines

The catalytically active version, exonuclease-dead (E84A), helicase-dead (K577M), and dually exonuclease and helicase-dead (E84A/K577M) versions of WRN were a gift from Raymond Monnat (Addgene plasmids # 46038, 46036, 46035, and 46037, respectively). The mis-sense mutant forms of WRN have been previously demonstrated to lack their indicated enzymatic activity^20^. The WRN sequence was cloned into a modified lentiviral expression vector, pLX_TRC209, under an EF1a promoter and modified to contain a neomycin selectable marker. Sanger sequencing of the vectors and genomic DNA after integration were performed to confirm sequence identity. Lentivirus was produced as described below and transduced into dually Cas9/Firefly-luciferase expressing KM12 to create stable ectopic-WRN-expressing cell lines.

### Lentiviral production

Lentiviral production was performed using HEK293Ts as described on the GPP portal (https://portals.broadinstitute.org/gpp/public/).

### sgRNAs

sgRNAs used in the validation studies were designed using the Broad Institute Genetic Perturbation Platforms sgRNA Designer (https://portals.broadinstitute.org/gpp/public/analysis-tools/sgrna-design). sgRNAs targeting *WRN* include sgWRN1 (target sequence GTAAATTGGAAAACCCACGG), sgWRN2 (target sequence (ATCCTGTGGAACATACCATG), sgWRN3 (target sequence GTAGCAGTAAGTGCAACGAT). sgRNA targeting the exon-intron junction (sgWRN-EIJ) were designed using the DESKGEN Cloud tool (https://www.deskgen.com/landing/cloud.html). The target sequence for sgWRN-EIJ is AGCACGTACATAAGCATCAG. Two negative controls targeting intergenic sites on chromosome 2 were utilized: sgCh2-2 (GGTGTGCGTATGAAGCAGTG) and sgCh2-4 (GCAGTGCTAACCTTGCATTG). Two pan-essential controls targeting *POLR2D* (AGAGACTGCTGAGGAGTCCA) and *c-Myc* (ACAACGTCTTGGAGCGCCAG) were used. sgRNAs were inserted in the pXPR_BRD003 lentiviral vector and inserts were verified by Sanger sequencing.

### shRNAs

shRNA targeting *WRN* were chosen from Project DRIVE. shRNA targeting *WRN* include shWRN1 (target sequence CAGCACTGCCAATGGTTCCAA) and shWRN2 (target sequence (GCCTTAACAGTCTGGTTAAAC). Positive pan-essential controls include shPSMD2 (target sequence CGCCAGTTAGCTCAATATCAT) and shRPS6 (target sequence CCGCCAGTATGTTGTAAGAAA). shRFP (target sequence CTCAGTTCCAGTACGGCTCCA) was used as a negative control.

### Immunoblotting

For immunoblotting, cells were lysed in RIPA buffer (Sigma) supplemented with cOmplete Protease Inhibitor Cocktail (Roche, 11697498001) and a Halt Phosphatase Inhibitor Cocktail (Thermo Fischer Scientific, 78428). Lysates were fractionated in 4–12% Bis-Tris gels (Invitrogen), which was then transferred to PVDF membranes (Immobilon-FL PVDF Millipore, IPFL00010) and blocked for an hour with Odyssey Blocking Buffer (PBS) (LI-COR Biosciences, 927-40000). Types of primary antibodies and the dilutions used for immunoblotting were as follows: anti-phospho-Chk2 [T68] (R&D Systems, AF1626, 1:400); anti-(total) Chk2 (Cell Signaling Technology, 3440, 1:1000); anti-γH2AX (Cell Signaling Technology, 9718, 1:1000); anti-GAPDH (Cell Signaling Technology, 5174, 1:1000); anti-MLH1 (Cell Signaling Technology, 3515, 1:1000); anti-MSH3 (BD Biosciences, 611390, 1:400); anti-WRN (Novus Biologicals, nb100–472, 1:1000). Subsequent steps of immunoblotting were conducted using Near Infrared (NIR) Western Blot Detection system (LI-COR Biosciences) as per manufacturer’s recommendations. Immunoblotting were performed three times. Shown are representative results from one experiment.

### Immunofluorescence

Immunofluorescence was conducted essentially as described previously^54^ (except for the double staining for WRN and fibrillarin; see below). Briefly, 2 days after lentiviral transduction, cells were seeded either on an 8-Well Lab-Tek Chamber Slide (Thermo Fisher Scientific, 177402) or on a 96-Well Clear Bottom Black Polystyrene Microplate (Thermo Fisher Scientific, 3904). The numbers of cells seeded per well were following: 1 × 10^6^ (5 × 10^5^), 1.2 × 10^6^ (6 × 10^5^), 1.6 × 10^6^ (8 × 10^5^), 6 × 10^5^ (3 × 10^5^), and 8 × 10^5^ (4 × 10^5^) cells for SW620, KM12, SW48, ES2, and OVK18 cells, respectively [numbers represent seeding densities for 8-well chamber (96-well plate)]. Cells were fixed and stained 2 days later. Micrographic images were acquired using either epifluorescence microscopy (for Figs. 3B, S4A, S4C and S4E) or confocal microscopy (for Figs. 4B, 4C, S5B, S5E, S5G and S6B), which were performed on an Axio Observer.Z1 microscope equipped with an Axiocam 506 mono camera and Apoptome.2 (Carl Zeiss) and a Zeiss LSM 700 laser scanning confocal system equipped with Axio Observer (Carl Zeiss), respectively. These confocal microscopy images represent maximum intensity projections of 5 consecutive planes with a step size of 0.08 μm. For image quantification, images were acquired using an Opera Phenix High-Content Screening System (PerkinElmer, HH14000000) and analyzed on Harmony High Content Imaging and Analysis Software (PerkinElmer, HH17000001). For phospho-p53-, p21-, γH2AX- and phospho-ATM-staining, signal intensities in the nucleus of at least 1000 cells/sample were scored on background-subtracted images and presented as box plots. The lower and upper limits of the box plot represent 25th and 75th percentiles, respectively and the bar in the middle of the box represent the median value. The low and high error bars represent 1st and 99th percentiles, respectively, Outliers are represented as dots. To score the patterns of nuclear staining, cells exhibiting mean signal intensity of 12000 or higher (for γH2AX) [20000 or higher for phospho-ATM] were first separated as cells with ‘pan-nuclear’ pattern of staining. For the rest of the cells, the number of foci within the nucleus was scored using a spot-detection program in the software. The relative abundance of cells exhibiting pan-nuclear staining and that of cells harboring specific number of foci were plotted. The number of nuclear foci observed in cells expressing the Apple-53BP1-trunc fluorescent marker were scored similarly.

IF for WRN and fibrillarin was performed as previously described^38^. Images were obtained via the Zeiss LSM510 Upright Confocal System for Fig. 4h. Weighted Pearson colocalization coefficients were calculated by obtaining Z-stacks of 5 representative high-powered fields at 63x magnification and scored via the Zeiss Zen Blue software. Significance was calculated by two-tailed Student’s *t*-test for MSI cell lines compared to lineage-matched MSS cells.

Types of primary antibodies and the dilutions used for immunofluorescence were as follows: anti-γH2AX (Millipore Sigma, 05-636, 1:400); anti-p21 (Santa Cruz Biotechnology, sc-6246, 1:100); anti-phospho ATM [S1981] (Millipore Sigma, 05-740, 1:200); anti-phospho Chk2 [T68] (R&D Systems, AF1626, 1:100); anti-fibrillarin (Abcam, ab5821, 1:500); anti-phospho p53 [S15] (Cell Signaling Technology, 9284, 1:100); anti-WRN (Sigma W0393, 1:200).

For all IF experiments except for the double staining for WRN and fibrillarin, the secondary antibody was Goat anti-rabbit (mouse) IgG, Alexa Fluor 488 (Thermo Fisher Scientific, A-11008 [A-11001]) was used at a 1:200 dilution. For WRN and fibrillarin IF, Goat anti-mouse IgG, Alexa Fluor 488 (Thermo Fisher Scientific, A-11001) and Goat anti-rabbit IgG, Alexa Fluor 555 (Thermo Fisher Scientific, A-21428) were used at 1:1000 dilution. Following secondary antibody treatment, the nuclei were counterstained with 4’,6-Diamidine-2’-phenylindole dihydrochloride (DAPI [Sigma, S9542]; 1 μg/ml). All IF experiments were performed twice. Shown are representative results from one experiment.

### Cell Viability Assay

Cas9-expressing versions of the following cell lines were seeded in 100 μL of media in 96-well plates (Corning 3904) excluding edge wells at the following densities: ES2 10^3^ cells/well, OVK18 1.5 × 10^3^ cells/well, SW620 2 × 10^3^ cells/well, KM12 2 × 10^3^ cells/well, SW837 2.5 × 10^3^ cells/well, SW48 2.5 × 10^3^ cells/well, GCIY 2 × 10^3^ cells/well, SNU1 at 1.5 × 10^3^ cells/well, Hec50B 1.75 × 10^3^ cells/well, SNGM 1.5 × 10^3^ cells/well. All cell lines except SNU1 were seeded the day prior to transduction. SNU1 (suspension line) was seeded on the day of transduction with 4 μg/ml polybrene. For the adherent cell lines, media was changed to media containing 4 μg/ml polybrene. Viral concentrations were pre-determined to achieve > 90% infection efficiency. Experiments were performed in triplicate by adding the appropriate volume of lentivirus to integrate vectors encoding the desired sgRNA and the plates were spun at 931 RCF for 2 hours at 30°C. The media was changed the next day and every 3 days thereafter. Cell viability was assayed using CellTiter-Glo (Promega G7572) at 33 μL per well. Luminescence was read using PerkinElmer EnVision 2105. Values were normalized to the average values from the negative control sgRNAs for each cell line. Experiments were performed three times. Shown are the triplicate results of one representative experiment. Two-tailed, type 3 Student’s *t*-test was performed to determine statistical significance, which was conducted on Microsoft Excel (Microsoft).

### Luciferase Competitive Growth Assay

Dual Cas9/Firefly-luciferase and Renilla-expressing cells were generated by transduction of Firefly-luciferase cDNA and Renilla-luciferase cDNA, respectively, in a pLX_TRC313 lentiviral expression vector containing a hygromycin-resistance gene. After selection, these two versions of a cell line were co-seeded in a 96 well plate at the following densities per well for each version: ES2 2 × 10^3^ cells/well, OVK18 3 × 10^3^ cells/well, SW620 4 × 10^3^ cells/well, KM12 4 × 10^3^ cells/well. The following day, transduction was performed in sextuplicate and media was changed to include puromycin the day after transduction. Cells were split and assayed with the Dual-Glo Luciferase Assay System (Promega) per the manufacturer’s recommendations every 3–4 days. Luminescence was determined by the Perkin Elmer EnVision 2105. Values are presented as the ratio of firefly to renilla luciferase luminescence signal per condition and normalized to the mean of the corresponding negative control sgRNAs for each cell line. Shown are the sextuplicate results of one representative experiment. Experiments were performed twice.

### Clonogenic assay

Cells were transduced with lentivirus expressing indicated shRNAs. 24 hours later, medium was replaced with 2 μg/mL puromycin containing medium. After 24 hours of puromycin selection, lentivirus-infected cells were trypsinized and reseeded onto a 24 well plate. The number of cells seeded per well were following: 3 × 10^3^, 4 × 10^3^, 6 × 10^3^, 1 × 10^4^, 8 × 10^3^ and 8 × 10^3^ cells for ES2, OVK18, SW620, SW837, KM12, and SW48 cells, respectively. Cells were subsequently propagated for 2 weeks in puromycin-free media, which was changed every 3 days. For crystal violet staining, cells were fixed with 10% formalin for 30 min at RT and subsequently stained with 250 μL/well of 0.1% crystal violet in 70% ethanol for 30 min at RT with constant shaking. To remove unbound crystal violet, cells were washed with DI water 3 times each for 5 min. Quantification was performed by extracting the crystal violet dye with 250μL of 10% acetic acid. 50 μL were transferred into a 96-well format in triplicate. The experiment was performed three times. Crystal violet absorbance was determined by the Perkin Elmer EnVision 2105. Shown are the results of one representative experiment with quantification demonstrating repeat measurements from a single experiment.

### Cell Cycle Analysis

Cas9-expressing cell lines were lentivirally transduced to deliver the desired sgRNAs. Media was changed the next day to allow for antibiotic selection. Four and seven days after the lentiviral transduction, cells were labeled with EdU, harvested, and stained as per the Click-iT™ Plus EdU Flow Cytometry Assay Kit recommendations. Stained cells were then examined with flow cytometry and results analyzed with FlowJo v10. Shown is a representative result of two independent experiments, each of which was conducted in triplicates. Statistical analysis of the proportion of cells in S-phase versus sgCh2-2 was calculated by the two-tailed Student’s *t*-test.

### Apoptosis Assay

Cas9-expressing cell lines were lentivirally transduced to deliver vectors encoding the desired sgRNAs. Media was changed the next day without antibiotic selection. Cells were split 4 days post transduction. Seven days post-transduction, cells were harvested to allow for Annexin V-FITC and Propidium Iodide staining. Stained cells were then examined with flow cytometry and results analyzed with FlowJo. Shown is a representative result of two independent experiments, each of which was conducted in triplicates. Significance is calculated for the sum of the proportions of cells in early apoptosis, late apoptosis and nonapoptotic death categories; vs. Ch2-2.

## Acknowledgments

The authors thank C.R. Boland, A. Goel, M. Koi, R.J. Monnat, and R. Weissleder for reagents; and A. Tang for graphical assistance. This work was funded by The Carlos Slim Foundation in Mexico through the Slim Initiative for Genomic Medicine and institutional funds from the Broad Institute. E.M.C. was supported by NIH Grant No. 2T32CA009172-42A1.

## Author Contributions

E.M.C., T.S., F.V. and A.J.B. initiated the project, designed, and supervised the research plan. J.M., M.G., Y.L. and Y.M. performed computational analysis of the CCLE and cancer dependency datasets under the supervision of D.E.R, J.S.B., G.G., T.R.G., A.T., F.V., and A.J.B. E.M.C., T.S., B.G., and J.S.M. performed experiments to validate the cancer dependency dataset findings with help from N.D., M.S., A.A., S.A.O., L.L., and A.G. RNA-seq analysis was performed by J.M. and M.I. J.S.M., T.S., and E.M.C. performed and analyzed the cell cycle and apoptosis assays. Immunoblots were performed by T.S., E.M.C., B.G., and J.S.M. Immunofluorescence were performed by T.S., E.M.C., J.B.L., J.L., E.A.R, and E.R. and analyzed by T.S., E.M.C., J.B.L., J.L., M.A., and A.D.A. E.M.C., T.S., J.F., F.V., and A.J.B wrote the manuscript. All the authors edited and approved the manuscript.

